# Modified RNA Extraction Methods to Eliminate Agarose Impurities in Precision-Cut Lung Slices

**DOI:** 10.64898/2026.02.16.705835

**Authors:** Reina Rangel, Sabina Anderson, Giuseppe DeIuliis, Edward P. Manning, Farida Ahangari, Abhay Pandit, Naftali Kaminski, Joan Marti-Munoz

## Abstract

Precision-cut lung slices (PCLS) have emerged as a powerful tool for studying the biology of viable human lung tissue. However, the presence of agarose impurities compromises RNA yield and integrity during the extraction process. We tested whether using an alternative Plant kit RNA extraction method to wash agarose impurities or pre-dissolving agarose from PCLS implementing a dissolving buffer for routine RNA isolation in gel-electrophoresis would improve RNA quantity, quality, and integrity. Our results show that RNA quantity and integrity are highly compromised when using a conventional method of RNA extraction. The plant kit and dissolution of agarose increased the RNA quantity to 0.42±0.11 and 0.65±0.17 µg/PCLS (measured by the Qubit) and integrity number to 6.60±0.59 and 9.13±0.39 (measured by the Bioanalyzer), respectively. The presence of impurities in conventional and Plant kit extractions misled to an overestimation of the RNA quantity and quality using the NanoDrop. The Plant kit and agarose dissolution showed a significant transcript integrity increase in GUSB (p<0.0001) and COL1A1 (p<0.05) expression, validating these methods over conventional extraction. We encourage laboratories applying PCLS experimentation to implement alternative methods to remove agarose impurities during RNA extraction, as well as to rely on sensitive quantitative techniques, such as the Qubit and Bioanalyzer, for RNA quantification and integrity measurements.

## INTRODUCTION

Precision-cut lung slices (PCLS) have emerged as a novel ex-vivo model to study biology in viable human lung tissue.(1, 2) The procedure involves low-melting agarose inflation to facilitate vibratome sectioning while maintaining the pulmonary microarchitecture. Despite these advantages, the presence of agarose impurities using conventional RNA extracting methods compromises quantity, quality and integrity.(3-5)

While methods to purify extracted RNA from cells encapsulated in agarose hydrogels (6, 7) or to dissolve agarose to isolate nucleic acids from gel-electrophoresis while preserving their integrity (8, 9) have been applied, these methods have yet to be implemented in PCLS.

In this study, we assessed whether using an alternative Plant kit extraction method to wash agarose impurities or pre-dissolving agarose from PCLS before RNA extraction by implementing a commercially available agarose-dissolving buffer improves RNA quantity, quality and integrity.

## METHODS

We generated PCLS from a recently deceased 74-years-old white male lung donor as previously described.(1, 2) Briefly, we inflated the lungs with warm 2 % (w/v) low-melting agarose solution, generated 12 mm ∅ cores at 4 °C, and sectioned 400 µm-thick PCLS in a VF-510-0Z vibratome (Precisionary).

We cultured PCLS in DMEM:F12K medium supplemented with 0.1% (w/v) bovine serum albumin and 1% (v/v) antibiotic-antimycotic with vehicle or fibrotic cocktail (FC)(10). After five days, we harvested PCLS for RNA extraction.

We monitored cell viability using Calcein and Ethidium homodimer-1 live/dead staining. We imaged the stained PCLS in a Leica Stellaris confocal microscope and quantified fluorescence on ImageJ.

We extracted RNA implementing three different methods: the conventional Qiagen miRNeasy micro kit (Agarose+), Qiagen RNeasy Plant mini kit to wash agarose impurities, or dissolution of agarose by immersing PCLS in ZymoResearch RNA agarose dissolving buffer for two minutes at room temperature, followed by RNA extraction using the Qiagen miRNeasy micro kit (Agarose-). (**Figure 1A**)

**Figure 1.**
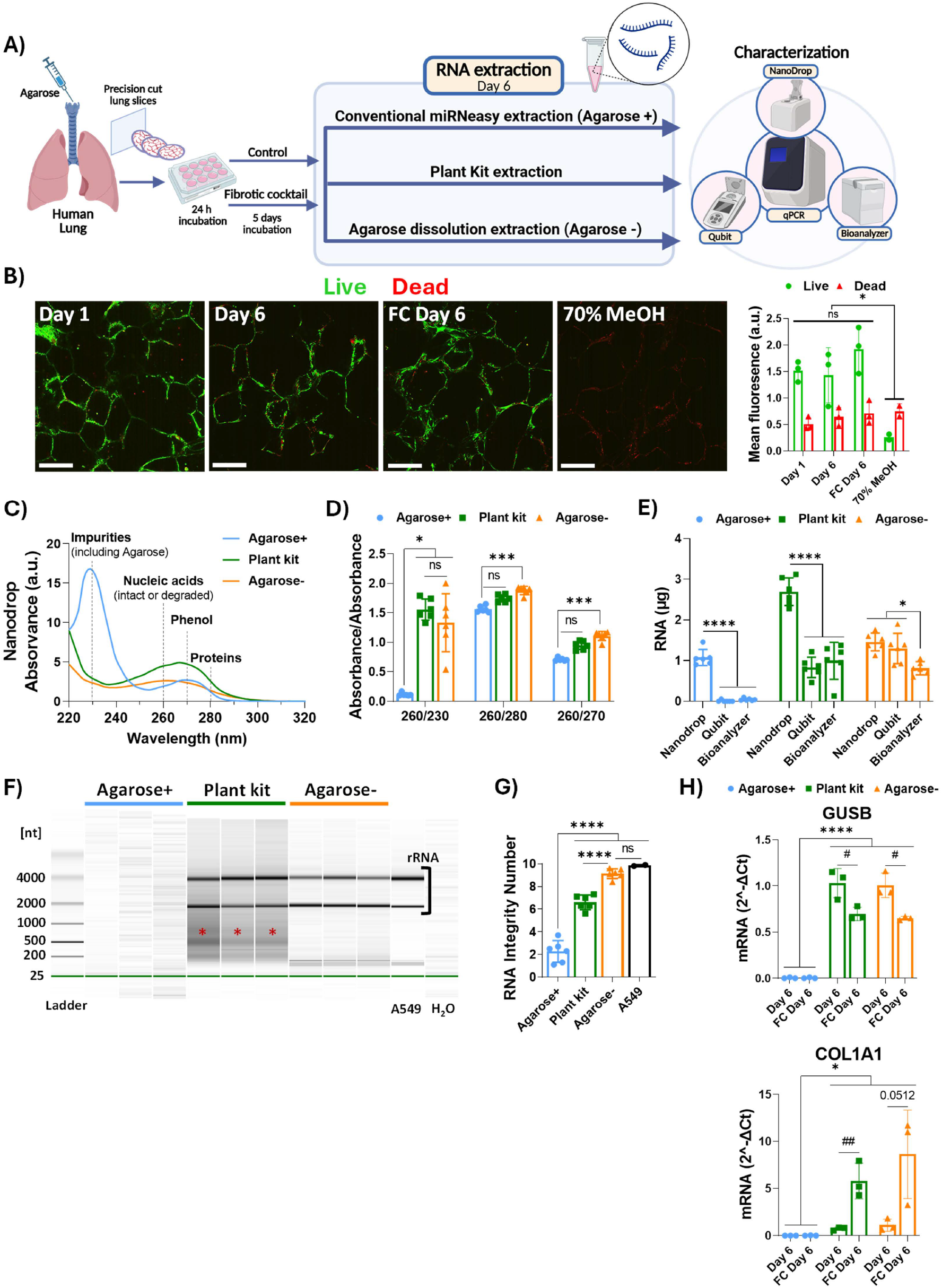
RNA extraction from PCLS using the conventional (Agarose+), Plant kit, and dissolution (Agarose-) methods. **A)** Workflow of the methodology. **B)** Representative live (green) and dead (red) confocal PCLS images and fluorescence quantification. PCLS incubated with 70 % MeOH for 30 min were included as dead control. n= 3 PCLS, scale bar = 350 µm. **C)** RNA and impurities absorbance for the different extracting methods measured by the NanoDrop and the **D)** 260/230, 260/280, 260/270 nm quality ratios. **E)** RNA quantification measured by the NanoDrop, Qubit, and Bioanalyzer. **F)** RNA electropherogram and **G)** integrity numbers (RIN) showing intact ribosomal (rRNA) and degraded (*) RNA measured by the Bioanalyzer. RNA extracted from A549 cells was included as positive control. **H)** GUSB and COL1A1 relative mRNA expression for the vehicle or fibrotic cocktail (FC) treatments measured by qPCR. n= 6 PCLS. Mean and SD. 2way or one-way ANOVA p ≤ 0.05(*), 0.001(***), 0.0001(****). Unpaired t test p ≤ 0.05(#), 0.01 (##).

We measured RNA quantity, quality and integrity in NanoDrop2000, Qubit2.0, and 2100 Bioanalyzer. We measured GUSB and COL1A1 gene expression by quantitative polymerase chain reaction (qPCR) in QuantStudio6Pro. Graphs and statistics were generated on GraphPad Prism 9.5.0 and illustrations were created on BioRender.

## RESULTS

Cells remained viable in PCLS when cultured for six days without or with fibrotic cocktail presence. No cells were viable when exposed to 70 % MeOH for 30 min (**Figure 1B**).

Conventional Agarose+ RNA extraction showed high presence of impurities absorbing at 230 nm, leading to a decrease in the 260/230 quality ratio of 0.12±0.02. Both the plant kit and agarose dissolution extractions efficiently removed the 230 nm impurities, suggesting efficient elimination of agarose. This led to an increase of the 260/230 quality ratio to 1.75±0.06 and 1.33±0.45, respectively (**Figure 1CD**).

In contrast, conventional Agarose+ and Plant kit extractions showed high presence of phenol contamination absorbing at 270 nm, leading to an overestimation of RNA quantity due to the overlap with nucleic acid absorbance at 260 nm. This also led to an overestimation of the 260/230 and 260/280 quality ratios.

RNA quantification using the Qubit and Bioanalyzer showed RNA yields below the detection limit for the conventional Agarose+ extraction and a ∼3-fold reduction for the Plant kit extraction (**Figure 1E**), confirming the inaccuracy of NanoDrop to quantify the RNA due to the presence of overlapping impurities. In contrast, agarose dissolution led to a clean extraction with the RNA absorbance peaking at 260 nm instead of 270 nm (**Figure 1C**), together with consistent values among the NanoDrop, Qubit, and Bioanalyzer measurements (**Figure 1E**).

Additionally, the Bioanalyzer showed lack of intact ribosomal RNA (rRNA) for the conventional Agarose+ extraction, indicating absence or high degradation of RNA (**Figure 1F**). The Plant kit and Agarose-dissolution extractions significantly (p<0.0001) improved RNA integrity. However, the Plant kit showed partially degraded RNA with RIN values of 6.60±0.59 compared to those of 9.13±0.39 for the Agarose-dissolution extraction.

The Plant kit and Agarose-dissolution extractions led to a significant increase in GUSB (p<0.0001) and COL1A1 (p<0.05) expression, indicating improved transcript integrity and validating these methods over conventional extraction (**Figure 1G, Table 1**). There were no differences in gene expression between both methods when implementing the more precise RNA quantification measured by the Qubit technique.

**Table 1.** GUSB and COL1A1 Cq values measured by the qPCR.

| **GUSB** |  |  |  |  |  |  |
| --- | --- | --- | --- | --- | --- | --- |
|  | **Agarose+** |  | **Plant kit** |  | **Agarose-** |  |
|  | **Day 6** | **FC** | **Day 6** | **FC** | **Day 6** | **FC** |
| **1** | 33.26 | 32.54 | 26.32 | 26.69 | 26.10 | 26.76 |
|  | 34.00 | 32.74 | 26.34 | 26.72 | 26.31 | 26.78 |
| **2** | 35.83 | UN | 25.92 | 26.45 | 26.01 | 26.70 |
|  | 36.96 | UN | 26.04 | 26.50 | 25.79 | 26.66 |
| **3** | UN | UN | 25.90 | 26.74 | 26.21 | 26.79 |
|  | UN | UN | 25.91 | 26.70 | 26.22 | 26.67 |
| **COL1A1** |  |  |  |  |  |  |
| **1** | 30.96 | 26.83 | 23.57 | 20.26 | 23.29 | 20.97 |
|  | 31.26 | 27.77 | 23.49 | 20.23 | 23.27 | 20.87 |
| **2** | 31.77 | UN | 22.94 | 19.68 | 21.80 | 19.17 |
|  | 31.80 | UN | 22.74 | 19.58 | 21.68 | 18.97 |
| **3** | 35.14 | UN | 22.89 | 20.52 | 22.97 | 19.21 |
|  | 35.59 | UN | 22.91 | 20.50 | 22.70 | 19.11 |

Cq values for 20 ng RNA in 10 $\mu$ L of Master Mix (TaqMan). Loaded RNA amounts measured by the Qubit. Two technical replicates per three biological replicates (n = 3 PCLS) per condition. FC = fibrotic cocktail, UN = Undetermined.

## DISCUSSION

The interpretation of PCLS experiments relies heavily on the analysis of gene expression. Therefore, obtaining pure, intact RNA is critical. In the past years, our laboratory has implemented PCLS experimentation to validate important findings.(11-13) However, we encountered inconsistencies when extracting RNA, leading to the invalidation and repetition of experiments.

Even if commercially available extraction kits are efficient at isolating RNA from biological tissues, they have not been optimized for PCLS.(3-5) Therefore, we validated alternative extraction methods.

Agarose is not efficiently removed by conventional extraction methods leading to large precipitates that likely retain RNA.(6) In this study, we showed that the quantity, quality, and integrity of RNA extracted from PCLS were highly compromised.

The RNeasy Plant kit was optimized to extract RNA from plant tissues which are rich in polysaccharides.(7) T. Ougura and co-workers identified this method as potentially beneficial to extract RNA from cells encapsulated in agarose hydrogels due to agarose being a polysaccharide.(6) Indeed, we observed a vast improvement in the reduction of agarose impurities in PCLS when using the Plant kit. However, phenol contamination was high, misleading to an overestimation of RNA quantity and quality.

On the other hand, molecular biologists have optimized the use of agarose dissolving buffers to purify nucleic acids from gel-electrophoresis.(8, 9) We showed that dissolving agarose from PCLS using an agarose dissolving buffer before RNA extraction rendered clean tissues for optimal extraction. Among the three extraction methods, agarose dissolution led to the cleanest and optimally intact RNA with consistent measurements across the different RNA quantification techniques. Removal of agarose impurities led to a significant increase in gene transcript integrity, supporting the importance of agarose removal in differential expression studies in PCLS.

In summary, we encourage laboratories implementing PCLS experimentation to apply alternative RNA extraction methods that efficiently remove agarose impurities, such as the ones presented in this study. We also encourage the use of the Qubit and Bioanalyzer techniques to precisely assess RNA quantity and integrity in PCLS experiments.

## Data availability statement

Approvals, detailed methodology, supplementary material and generated data can be accessed on https://data.mendeley.com/datasets/r6ypkfg2pc/1

Rangel, Reina; Anderson, Sabina; DeIuliis, Giuseppe; Manning, Edward P.; Ahangari, Farida; Pandit, Abhay; Kaminski, Naftali; Marti Munoz, Joan (2026), “Modified RNA Extraction Methods to Eliminate Agarose Impurities in Precision Cut Lung Slices_supplementary”, Mendeley Data, V1, doi: 10.17632/r6ypkfg2pc.1

## Acknowledgments

We thank the National Disease Research Interface (NDRI) for providing the lung sample.

## Contributors

RR, NK and JMM developed and supervised the study. AP, NK, and JMM secured funding and were involved in the conceptualization. GD, EM, FA, and NK managed ethical approval and organized the shipment of the lung. RR, SA, and JMM generated PCLS and performed experiments. RR optimized the Plant Kit extraction. JMM optimized the agarose dissolution extraction. RR, NK, and JMM analyzed the data. JMM developed the manuscript figures. RR, NK, and JMM contributed to the preparation of the manuscript. All authors have reviewed and edited the final manuscript. JMM is the guarantor of the manuscript.

## Ethics approval

This study was approved by the Yale University Institutional Review Board (2000022618).

